# Low-cost genomics enable high-throughput isolate screening and strain-level microbiome profiling

**DOI:** 10.1101/2022.04.11.487950

**Authors:** Jon G. Sanders, Weiwei Yan, Andrew H. Moeller

## Abstract

Earth’s environments harbor complex consortia of microbial lineages that affect processes ranging from host health to biogeochemical cycles. However, understanding the evolution and function of these microbiota has been limited by an inability to isolate individual microbial constituents and assemble their complete genomes in a high-throughput manner. Here, we present a workflow for bacterial isolation and whole-genome sequencing from complex microbiota using open-source labware and the OpenTrons automated liquid handling robotics platform. Our approach circumvents the need for isolate screening (e.g., through 16S rDNA sequencing or mass spectrometry analyses) by reducing the costs of genome-sequencing to ~$10 per bacterium. Applying the workflow, we quantified genomic diversity within 45 bacterial species in the chimpanzee gut microbiota. Results revealed hotspots of recombination in bacterial genomes and elevated transmission of plasmids between distantly related bacterial species within individual chimpanzee hosts. This study develops and applies an approach for high-throughput bacterial isolation and genome sequencing, enabling population genetic analyses of bacterial strains within complex communities not currently possible with metagenomic data alone.

## Introduction

Metagenomes are complex mixtures of organisms, with dozens to hundreds of microbial species sharing genes both through ancestry with closely related strains, as well as through horizontal transfer to distantly related lineages [1–3]. Understanding how genetic variation arises and changes within these communities is critical if we hope to develop useful models of their evolution [4].

But despite the tremendous advances in sequencing technology in the past decades, the paired phenomena of within-species strain diversity and between-species horizontal gene transfer still present a challenge to assessing the genetic structure of populations within diverse metagenomes like the mammalian gut. Community metagenome sequencing can rapidly generate massive quantities of data from a microbiome, but with only limited ability to link genetic changes within the same genome or in populations of closely related cells [5,6]. Mobile DNA elements, especially plasmids, are even more difficult to place in a metagenomic context [7–9].

In principle, cultivation offers a much more robust way to explore genomic variation within populations. By confidently drawing cellular bounds around genes, isolation represents a gold standard for describing genomic diversity and a necessary prerequisite for empirically demonstrating the functional consequences of such variation. As a consequence, cultivation has seen renewed interest, even earning its own ‘omics’ appellation: high throughput approaches to culturing, or ‘culturomics’ [10]. However, such high-throughput approaches typically require enormous investments in capital equipment and labor [10–13], putting them out of reach for many researchers. This is especially true for those studying non-model systems where the bulk of unstudied microbial diversity is likely to be found. While advances in miniaturization and microfluidic technologies may one day permit rapid high-throughput cultivation from diverse environments [14,15], such approaches are not yet widely available.

The recent availability of distributed, open-source laboratory automation and distributed manufacturing technologies suggests an alternative approach: adapting high-throughput cultivation techniques to relatively inexpensive commercial and in-house-manufactured equipment. In combination with the extremely low per-base cost of modern sequencing, such an approach offers the potential to realize much of the benefits of capital-intensive conventional high-throughput culturing and sequencing pipelines at a fraction of the required investment.

Motivated by our desire to explore genomic evolution in the microbial populations associated with natural mammalian gut microbiomes, we set out to design an inexpensive end-to-end high-throughput cultivation and genome sequencing protocol that could be easily replicated with a minimum of capital expenditure. We developed protocols, 3D-printed custom labware, and analysis pipelines to enable cost effective high-throughput cultivation and whole-genome sequencing of natural gut microbiota. These methods allowed us to circumvent traditional 16S rDNA- or mass spectrometry-based screening approaches, instead using full-genome sequencing to identify all cultivated isolates. Moreover, this approach enabled the generation and assembly of thousands of bacterial genomes from the chimpanzee gut microbiota rapidly and at low cost relative to existing approaches. Results revealed substantial variation in the distribution of strain-level diversity among chimpanzee hosts, and, importantly, allowed us to link putative plasmids to their specific bacterial hosts across chimpanzee individuals, populations, and subspecies.

## Methods

To accomplish our goals of maximum isolate genome throughput with minimal capital and labor costs, we developed a workflow based around the OpenTrons OT-2 robotic liquid handling platform (Fig. 1). This instrument allows for repeatable automation of many protocols, while costing less than $10,000 as configured. Where possible, we took advantage of previously-published low-cost molecular biology protocols, adapting them for automation on the OpenTrons platform. All the protocols described here are available at https://github.com/tanaes/Moeller_Opentrons_protocol_library. In addition, we wrote extensions to the OpenTrons Protocol API to improve certain aspects of instrument behavior, especially relating to use with magnetic bead protocols. An installable library of these extensions is available at https://github.com/tanaes/opentrons_functions.

**Figure 1.**
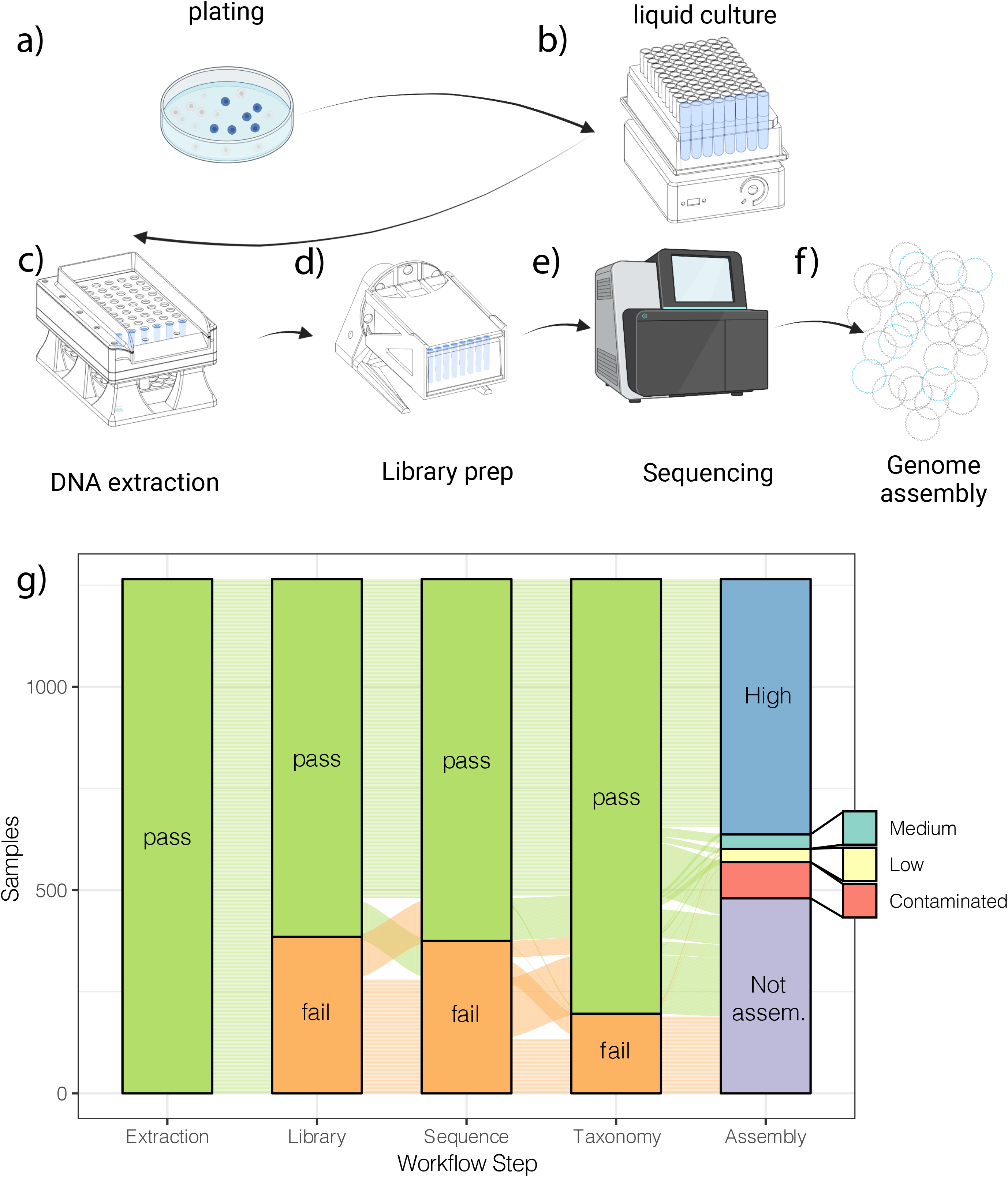
Illustration of isolate genome screening workflow, highlighting 3D-printed labware. a) Dilution plating on standard media. b) Liquid culture in 1.8 mL strip tubes, using 3D-printed compact plate shaker to enhance nutrient and gas mixing. c) DNA extraction on Opentrons OT-2 platform, using 3D-printed bead dispenser to aliquot lysis beads directly into liquid culture tubes. d) Library prep on Opentrons OT-2 platform, using 3D-printed plate rotator to enhance efficiency of DNA binding to magnetic beads. e) DNA sequencing on Illumina platform. f) Genome analysis and assembly. g) Results from initial rounds of screening, showing samples passing certain QC thresholds at each stage (extraction: 0.1 ng/μL DNA concentration; Library prep: 0.5 ng/μL DNA concentration; Sequencing: 25 Mbp sequence yield; Taxonomy: taxonomy assigned by Sourmash; Assembly: High, Medium, and Low-quality assemblies), Contaminated assemblies, and unassembled samples. Colored lines connect the same sample through each stage of the chart. Note that even many samples with low DNA extraction concentrations often yielded sufficient sequence data for taxonomic assignment.

For some protocols, we found that there were key steps that would require laboratory apparatus that were either not available for commercial purchase, uncommon in a typical molecular biology lab, or would require substantial investment. For these steps, we designed our own versions suitable for rapid manufacture with 3D printers and/or laser cutters, and using inexpensive commodity electronic components. These apparatus were designed using Fusion360 CAD software (Autodesk, Inc.). Full source files and component lists can be found at https://github.com/CUMoellerLab/Labware. All data in this paper were generated using versions of the 3D-printed apparatus described below, rather than commercially purchased alternatives.

### Sample collection, storage, and metagenome sequencing

The samples used in this study comprise 10 fecal specimens collected from wild Chimpanzees and Bonobos between July, 2003 and August, 2014 (Table S1); samples and sequences have been described previously [16]. Briefly, fecal samples were stored in RNALater at −80°C prior to use. Previous work has shown that RNLater preservation acts as a selective agent on microbial diversity within mammalian fecal samples [16], thereby allowing selected cultivation of the subset of the microbiota that remains viable. Here, using these RNALater-preserved samples enabled us to conduct high-powered analyses of intraspecific variation within a few dozen bacterial clades of interest, including *Clostridum* and *Bacilli*.

For metagenome sequencing, samples were centrifuged and approximately 50 mg of material removed from the pellet for DNA extraction. We extracted metagenomic DNA from pellets using the Qiagen PowerSoil extraction kit. Libraries were generated from metagenomic DNA using the “Illumina Equivalent” library prep method at the Cornell Biotechnology Research Center, and pooled libraries sequenced using an Illumina NovaSeq instrument at the UC Davis Sequencing Center.

### Cultivation

Twelve fecal samples were selected for cultivation. To sample as much diversity as possible, we used several different media for cultivation: Yeast Casitone Fatty Acids (YCFA), YCFA+Starch, *Bifidobacterium* selective media (BSM), Brain heart infusion-supplemented (BHIS), and *Bacteroides* Bile Esculin (BBE) (Table S2). Recipes for all media were derived from [16]. For each sample-by-medium combination, 100 μl of fecal material suspended in RNAlater was plated in an anaerobic chamber (Coy brand) on solid media. Plates were incubated at 37°C for five days in an anaerobic (5% hydrogen, 5% carbon dioxide and 90% nitrogen) chamber (Coy Lab Products Inc).

Liquid culture of picked colonies represented a potential throughput bottleneck, especially if isolates were cultured in conventional glass test tubes. To increase throughput, we instead grew colonies in 1.2 mL 96-place strip tube racks, which have the footprint and well spacing necessary for processing on the OpenTrons liquid handler. Individual colonies were picked from plates into 900 μL of liquid media (Table S2) using a sterile wooden toothpick. Then, plates were incubated at 37° in the anaerobic chamber for four days.

To improve growth in liquid culture for cells that might benefit from increased waste gas diffusion or nutrient distribution, we designed small single-plate orbital shakers to fit inside our anaerobic incubator (Figure 1a). Adapting an existing open-source design (https://learn.adafruit.com/crickit-lab-shaker/3d-printing), we simplified the electronic components, relocated all connections and controls to the front of the apparatus to facilitate use within the incubator, and changed it to use 5V USB input for power, allowing us to use a single USB charger to power 7 individual shakers within the incubator.

Following incubation, 300 μL of media per tube was transferred to a clean deep-well plate and cells pelleted in a centrifuge at 16,000 *g*. After removal of supernatant, cells were resuspended in glycerol buffer and stored at −80°C for future use.

### DNA extraction

Kit-based DNA extraction protocols typically cost between $3 and $5 per sample. For 16S amplicon-based screening, this step can sometimes be omitted with a chemical lysis prior to amplification. For whole-genome screening, though, we judged that the added complexity of a DNA extraction step was necessary. To reduce costs, we adapted the magnetic bead-based extraction methodology from [17], which uses laboratory-made reagents and either purchased or lab-made magnetic beads, for use on the OpenTrons platform.

For cell lysis, we chose to use beadbeating to ensure lysis of a broad range of bacterial cell types. We designed a 3D-printed and laser-cut loading system to precisely load 0.2 mm glass beads directly into the 96-well strip-tube plates (Figure 1b) after pelleting cells and removing liquid media. After bead loading, 800 μL of guanidine HCL lysis buffer was added to the tubes, and they were capped and shaken on an Omni Bead Ruptor Elite at 6.5 m/s for 40 seconds. The tubes were then spun down on a centrifuge at 400xg for 5 minutes, decapped, and then moved to the OpenTrons instrument for the remainder of the extraction. The detailed OpenTrons extraction protocol can be found in the project repository linked above. Briefly, the robot transfers 600 μL of lysate to a new plate, adds magnetic beads in a PEG-based binding buffer, and then goes through a series of magnetic binding and wash steps before eluting the extracted DNA in nuclease-free water.

We found extraction efficiency was greatly improved by gently agitating magnetic beads during the initial binding step. To accomplish this, we designed a 3D-printed rotator (Figure 1c) with attachments for holding 96-well plates or microcentrifuge tubes. After transferring lysate and adding beads and binding buffer on the liquid handler, we programmed in a pause to allow the user to remove the plate, seal it, and place it on the rotator for 10 minutes. Following this step, the plate was unsealed and returned to the liquid handler for the remainder of the protocol.

Extracted DNA was quantified in 384-well plates using a reduced-volume version of the QuantiFluor (Promega) fluorescence-based assay. Four 96-well plates (each the output from a single extraction protocol) were tested in each assay, using an OpenTrons protocol for sample transfer and a Tecan Infinite M200 plate reader for quantification.

### Library prep and sequencing

To inexpensively generate sequencing libraries from thousands of DNA extractions, we adapted the Hackflex library prep protocol [18] to the OpenTrons liquid handler. Briefly, this protocol dilutes key reagents from the Illumina Library Prep protocol to stretch a single kit across more samples. Our adaptation of the protocol changes some reagent quantities to better fit the constraints of the OpenTrons format; for details, see the full protocol in the project repository linked above.

For the libraries presented here, we used barcoded library amplification primers provided by the Cornell Biotechnology Resource Center. Initially, these shared a single i5 index per library plate, with unique i7 primers per sample. For later libraries, we switched to unique dual indexed (UDI) primers, with 96 unique i5 and i7 primers per plate. To facilitate multiplexing across library prep plates with UDIs, we created a version of the protocol to cycle column matches between i5 and i7 primer plates, allowing up to 12 library plates to be multiplexed without repeating an index combination. Libraries were amplified using 17 cycles of PCR prior to bead-based dual-sided size selection and final elution.

Final libraries were quantified by QuantiFluor (Promega) in 96 well plates, then pooled according to the following algorithm: the volume necessary to transfer 5 ng of library DNA was calculated; for samples requiring more volume than this to reach 5 ng transferred (likely failed libraries), 1 μL was transferred; for samples requiring less volume than this, 0.5 μL was transferred. Per-plate pools were combined and concentrated using magnetic beads and then provided to the Cornell Biotechnology Resource Center for sequencing on an Illumina NextSeq 500 instrument. Two separate sequence runs were performed, combining 13 and 10 library prep plates, respectively.

### Sequence analysis

Isolate sequences were processed using the Bactopia pipeline [19], which does sequence trimming and QC with FastQC (https://www.bioinformatics.babraham.ac.uk/projects/fastqc/), assembled sequences with Shovill (https://github.com/tseemann/shovill) and SKESA [20], performs assembly quality checking with CheckM [21], and gene annotations with Prokka [22]. Additionally, we used Bactopia-Tools to perform pangenome analysis with Roary [23] and recombination prediction with ClonalFrameML [24]. To create a phylogeny of isolates, assembled genomes predicted to be less than 5% contaminated with CheckM were processed using PhyloPhlAn2 [25] using the Amphora2 marker set [26] and the “Fast / High Diversity” default settings. To estimate abundances of isolates in original samples, we used CoverM (https://github.com/wwood/CoverM) to calculate coverage for each isolate genome in each of the available chimpanzee metagenomes. To estimate genome-wide recombination rates, we used ClonalFrame ML using 50,000 bp sliding windows and default settings.

We next identified plasmids within each isolate genome assembly to assess the distribution of these mobile elements among bacterial and chimpanzee hosts. We designated any contig predicted to be a complete circular element by Shovill as a putative plasmid. To estimate plasmid sharing between isolate genomes, we aligned each contig identified as a putative plasmid against each other in an all-by-all manner using BLASTn [27]. Contigs that aligned to one another across 98% of their length and had greater than 90% sequence identity were considered to be in the same plasmid ‘group.’ A bipartite network linking isolate genomes to plasmid groups was constructed in Python using the NetworkX library [https://networkx.org], and imported for visualization in CytoScape [28].

## Results

For the purposes of validating the workflow, we carried every sample from DNA extraction through to sequencing. Even if, for example, an isolate failed to grow during liquid culture, we did not exclude it from downstream steps. This enabled us to determine appropriate exclusion criteria for future use.

In total, we picked, grew in liquid culture, extracted DNA from, and sequenced 1911 isolates. Of these, 1295 yielded extractions with DNA concentrations above 0.1 ng/μL; 1070 yielded library concentrations ≥ 0.5 ng/μL; 1066 yielded ≥ 25 Mbp of sequence; and 761 yielded high-quality assemblies (> 90% complete and < 5% contaminated), 52 medium-quality assemblies (>50% complete and < 5% contaminated), and 58 low-quality assemblies (≤50% complete and < 5% contaminated) (Figure 1e). In total, 113 of the sequenced libraries gave assemblies that appeared to be contaminated based on CheckM results, indicating that around 10% of picked colonies may have not in fact been single clones.

The primary point of failure in the workflow appeared to be the liquid culture phase: only 68% of isolates yielded DNA concentrations above 0.1 ng/μL, and 32% above 1 ng/μL (Figure S1). Low turbidity of many tubes after incubation was consistent with either slow or no growth in liquid media for many of the colonies transferred from plated media. Initial DNA concentration was a good predictor of subsequent performance: 895 of the 1070 libraries with concentrations ≥ 0.5 ng/μL came from DNA extractions with concentrations above 0.1 ng/μL. 747 of the 761 high-quality assemblies (98%) came from samples with library concentrations above 0.5 ng/μL (Figure S2).

### Isolate diversity and distribution

Of the fully-assembled isolate genomes, 734 were classified successfully with GTDB-Tk. All 734 were classified as Firmicutes, with most (572) belonging to the Bacilli and 162 to the Clostridia. Together, these accounted for 9 unique taxonomic assignments at the order level, 13 at the level of family, and 31 at the genus level; all 734 genomes were assigned to a genus. Most genomes (603/734) were also assigned to a species. By far the most common genus among the assembled genomes was Streptococcus (348), followed by Enterococcus ‘D’ group (76), Staphylococcus (53), Claustridium ‘P’ group (46), and Blautia ‘A’ group (42). Taxonomic classifications, assembly statistics, and other metadata for all isolates are presented in Additional File 1.

Sourmash was able to classify more of the samples, with 1531 being classified to at least the phylum level. 983 were classified to order, 909 to family, 894 to genus, and 874 to species level. These classifications were highly consistent with the full-genome taxonomies, with 98.7% matching at the phylum and class levels, 97.2% matching at the order and family levels, 97.0% matching at the genus level, and 94.3% matching at the species level. Phylogenetic reconstruction using concatenated marker gene sequences was also largely concordant with taxonomic assignment (Figure 2).

**Figure 2.**
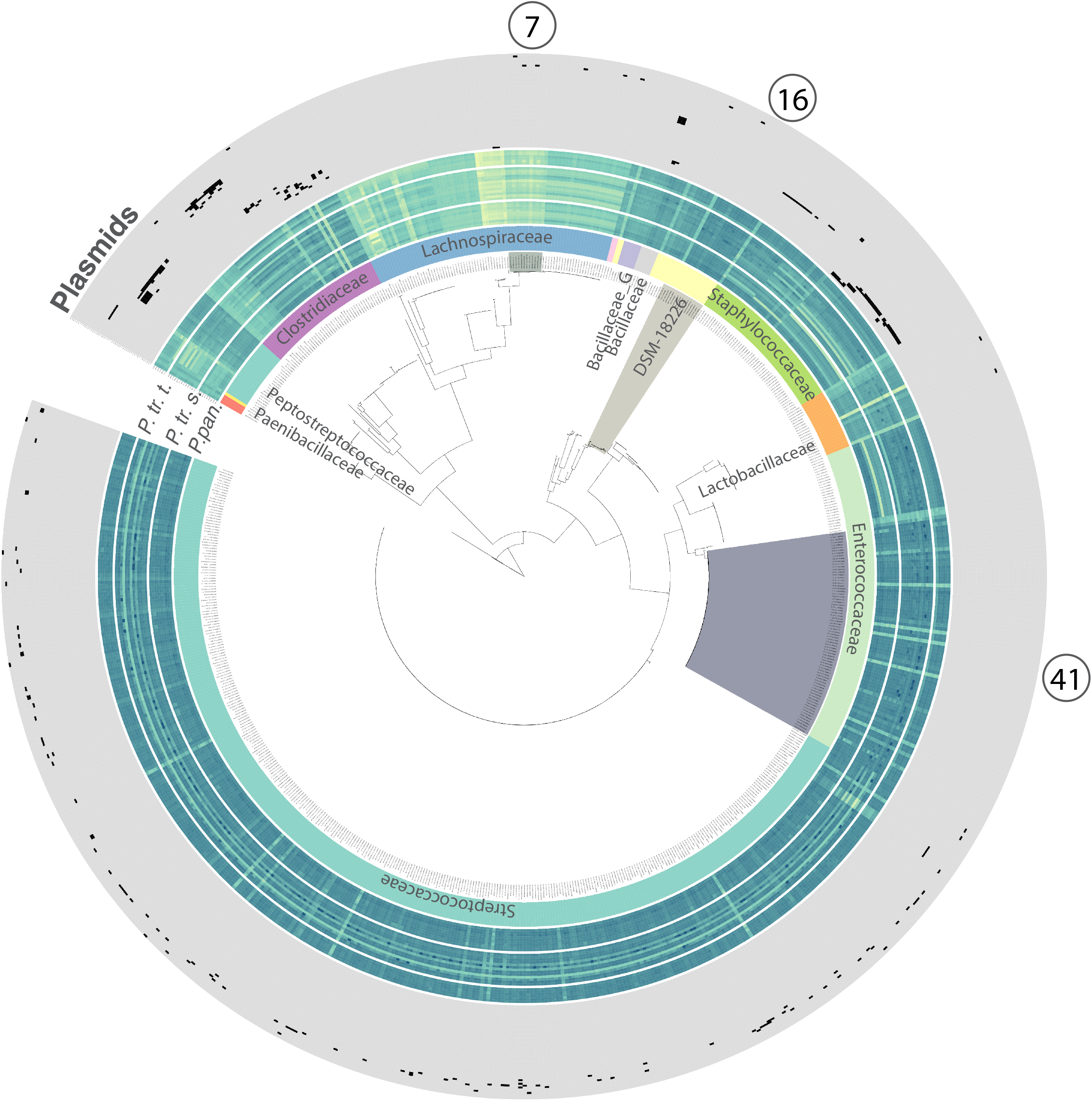
Multilocus phylogenetic reconstruction from 706 isolate assemblies using Phylophlan and the Amphora2 universal single-copy marker gene set. Colors and labels on the inner ring indicate family-level taxonomic assignment from GTDB-tk. Heatmaps in middle rings indicate log10 estimated coverage per isolate genome from CoverM within metagenomes of wild *P. paniscus*, *P. troglodytes schweinfurthii*, and *Pan troglodytes troglodytes*. Each grey outer ring indicates presence (black) or absence (grey) of a putative plasmid within bacterial isolates.

Mapping metagenomic reads sequenced directly from wild chimp fecal samples against the assembled isolate genomes supported an origin from those samples. A mean of 2.97% (SD 1.12%) of metagenomic reads mapped to the isolate genome assemblies.

### Isolate population genetics

Assembling individual isolate genomes also allowed us to explore the variation in within-species diversity that would have been hidden by 16S rRNA gene-based screening. Using dRep [29], we clustered the assembled isolates into 45 clusters sharing genome-wide estimated Average Nucleotide Identity of > 95%. All-by-all ANI comparisons within these clusters indicated differences in within-cluster diversity among clusters (Figure 3a, Figure S3).

**Figure 3.**
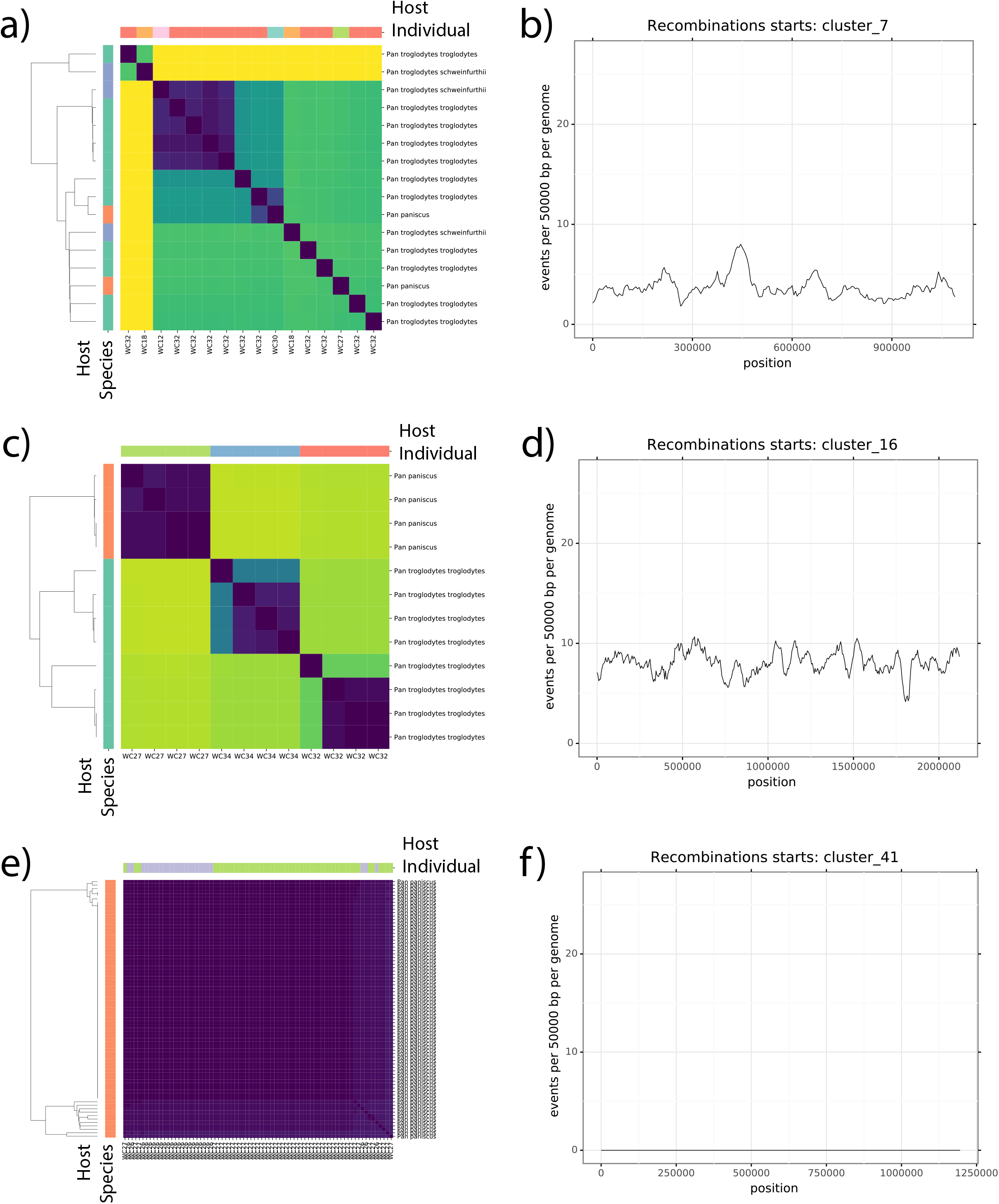
Differences in genome diversity observed among three example clusters of closely related strains. a, c, and e) Pairwise Average Nucleotide Diversity among strains within each cluster show different patterns of within-‘species’ diversity revealed by whole-genome screening. Heatmap color values indicate log pairwise nucleotide diversity between each pair of isolates in a cluster. Color bars at left and annotations at right show host species identity of the sample from which the isolate was recovered. Color bars at top show the host individual. b, d, and f) 50kbp sliding window of recombination events inferred by ClonalFrameML for strain clusters 7, 16, and 41, respectively.

In addition, the assembly of whole-genomes from isolates without dereplication (e.g., based on 16S rDNA similarity) enabled us to estimate genome-wide recombination rates within bacterial species (i.e., 95% ANI clusters). These analyses revealed non-uniform rates of recombination among bacterial lineages and across loci within lineages (Figure 3b, Figure S4).

### Putative plasmid diversity and distribution

375 isolate assemblies yielded a total of 622 circular contigs, which we are considering as putative plasmids. Most of these circular contigs were less than 50 kbp in length, with a mean length of 12,656 bp and a median of 5048 bp. Circularized contigs displayed significantly higher mean coverage than the overall assembly (3.89x higher than background; Figure S5), consistent with their annotation as putative plasmids. In addition, most of these circular contigs (143) were unique in our dataset, meaning there were no other circular contigs assembled that were at least 80% identical across at least 90% of their total length. The remaining contigs could be grouped into 81 unique types, shared between 2 to 63 different isolate assemblies (Figure 4).

**Figure 4.**
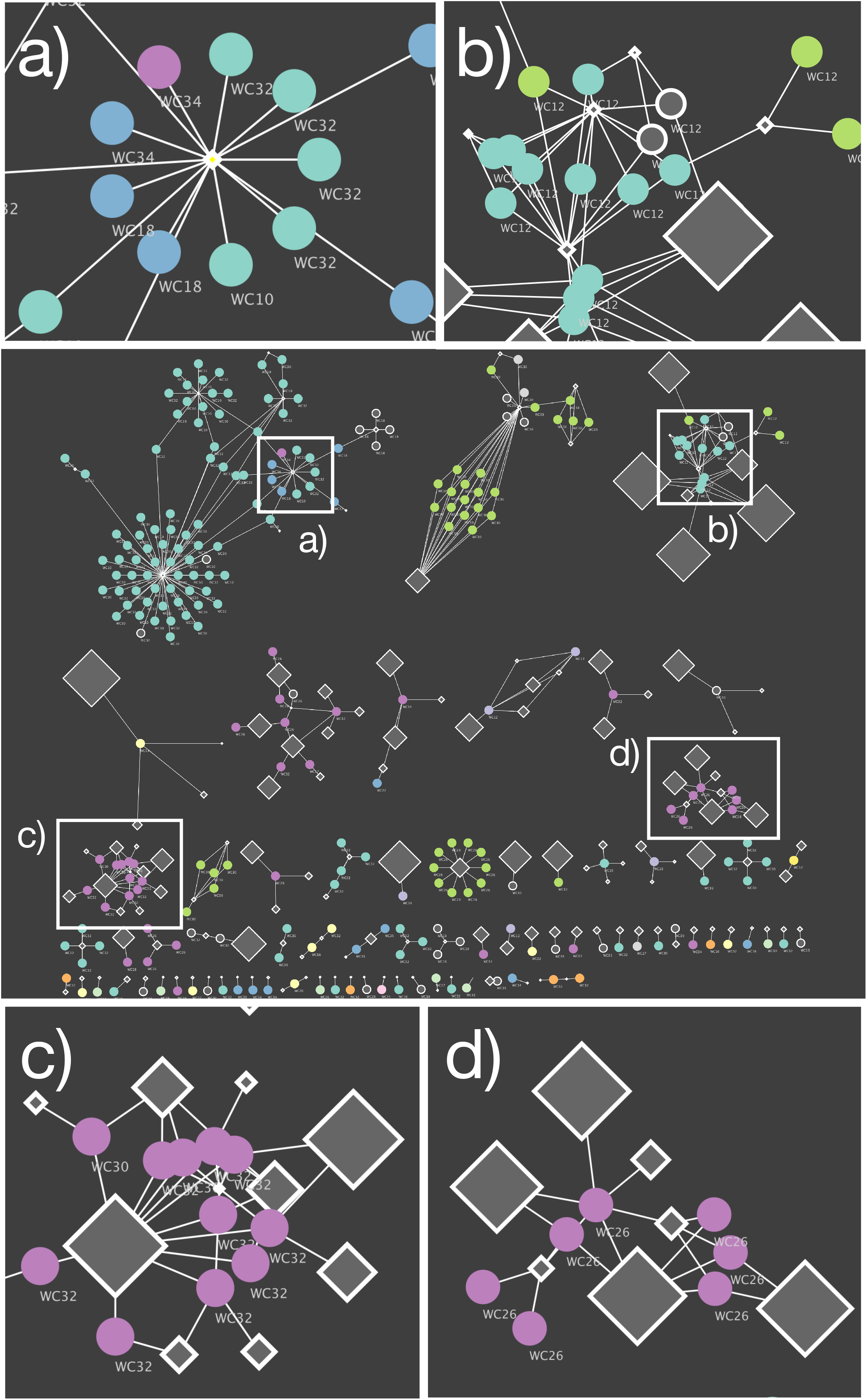
Plasmid sharing among distantly related bacteria within hosts. Bipartite networks illustrate relationships between circularized contigs (diamond-shaped nodes) and the isolates (circles) from which they were assembled. White edges connect circular contigs with each isolate from which they were recovered. Contig-node size is scaled approximately with the log of the total length of the contig. Circular isolate-nodes are colored according to the taxonomic class assigned to the isolate, and annotated with the individual from which they were cultured. Insets highlight specific regions of interest. a) A small circular contig recovered from several different unrelated bacterial taxa found several different host individuals. b) A sub-network of circular contigs that appears to be shared among different and sometimes unrelated isolates but all from the same host individual, suggesting the exchange of this genetic material among cells within the lifetime of the host. c and d) Sub networks of circular contigs that appear to be shared primarily among related bacteria from the same host individual.

Consistent with phylogenetic and geographic barriers to exchange of plasmids, we found that circular contigs were most often shared between genomes of related taxa found within the same host individual (Figure 4c, d). In many cases, several unique plasmids formed clusters within groups of related bacteria from the same host individual (Figure 4d). In other cases, though, the same circular contigs could be found in genomes from multiple distantly related bacterial taxa, either within the same host individual (e.g. Figure 4b), or even across many different host individuals (Figure 4a).

### Protocol cost estimates

Costs are difficult to estimate and communicate accurately, as purchasing prices and available equipment vary widely among laboratories. However, as one of the primary motivations of this manuscript is to make high-throughput isolate genome sequencing accessible to as many researchers as possible, we give our best estimates for both our required capital investment and per-sample consumable costs (Table S3) as a point of reference.

Our laboratory already had basic molecular biology equipment, including PCR machines, centrifuges, manual pipettes, and access to a fluorescence plate reader and bead beater. Additional capital expenses required for this protocol included an OpenTrons OT-2 robot with 2 multi-channel pipettes and a magnetic plate expansion module, a strip tube bead beater adapter, and materials costs for the 3D-printed labware; in total, capital expenses amounted to approximately $13,000.

We estimate a per-sample consumables cost of around $7.50. Of this, liquid culture and DNA extraction accounts for about $1.50; library preparation around $3; and sequencing around $3. Even with the 36% success rate we observed here, with no culling of failed samples prior to sequencing, this equates to around $20 per high-quality genome assembly.

## Discussion

We developed a workflow for high-throughput bacterial isolation and low-cost genome sequencing from complex microbiota. Our workflow makes use of custom-designed 3D-printed labware, the relatively inexpensive OpenTrons liquid handling platform, and recently developed methods for Illumina library preparation using highly diluted reagents. Together, this combination of methods allowed the isolation, library preparation, and whole-genome sequencing of hundreds of isolates in parallel for costs of ~$10 per bacterium. Importantly, by reducing per-isolate whole-genome sequencing costs substantially, our workflow alleviates the need for 16S rDNA- or mass spectrometry-based approaches for dereplicating bacterial strains prior to whole-genome sequencing. Of the bacterial isolates that grew in liquid culture and yielded appreciable DNA concentrations (>0.1 ng/uL) post-extraction (i.e., ‘Pass’ in Columns 1 and 2 in Figure 1g), >80% yielded Hackflex libraries, nearly all of which in yielded genome drafts upon sequencing (Figure 1).

Compared to metagenome-based approaches, isolation of individual bacteria affords the opportunity to sequence whole genomes and their associated plasmids without reliance on statistical inferences. We demonstrate the utility of this approach by isolating and profiling strain-level bacterial diversity in the chimpanzee gut microbiota. Results enabled the examination of intraspecific patterns of bacterial polymorphism, the estimation of genome-wide recombination rates without reliance on assumptions regarding phasing, and the identification of plasmids associated with specific bacterial and chimpanzee hosts. Thus, isolate genomics enables population-genetic analyses of bacterial species that remain difficult with shotgun-sequencing data alone.

Analyses of plasmid distributions among bacterial and chimpanzee hosts revealed several instances in which distantly related bacteria shared plasmids within—but not between—chimpanzee hosts, populations, and subspecies. These results support the view that co-occurrence within a shared environment increases rates of horizontal gene transfer (HGT), as has been indicated previously by metagenomic surveys [13,30,31]. Interestingly, our results suggest that the signatures of these HGT events are detectable even within the lifespan of individual hosts (Figure 4).

Our workflow has several advantages and disadvantages relative to existing approaches for high-throughput bacterial isolation and whole-genome sequencing. One major advantage of our workflow is its simplicity, as it relies on standard microbiological and molecular biology approaches and is fully automated on the OpenTrons platform. For example, relative to microfluidics-based isolation [32,33] or single-cell genome sequencing approaches [34,35], our method is readily applicable by labs without the need for capital intensive specialized equipment. The equipment costs necessary to execute our full protocol are also dramatically lower than for a number of previously-developed high-throughput genome sequencing workflows that achieve low marginal costs using expensive robotics capable of handling nanoliter-scale reaction volumes [36,37]. Similarly, while Hi-C based approaches also have the ability to link plasmids with their host bacteria, these methods rely on labor intensive protocols that crosslink chromatin with formaldehyde, then digested, and re-ligated to isolate covalently linked DNA fragments [34]. Moreover, both droplet and Hi-C approaches typically capture only a fraction of the genome, and they in general do not allow for the retention of isolated cultures for further experimental study. In contrast, a weakness relative to single cell and Hi-C approaches is that our workflow can only interrogate bacteria that are cultivable and amenable to isolation.

The data we report here represent the first two complete full-scale sequencing runs from this protocol, and there are still opportunities for improvement. First, although an overall hit rate of ~ 2 high-quality genome assemblies per picked 5 clones still represents substantial cost savings relative to more conventional methods, we are confident it could be improved. Indeed, as we have improved our anaerobic liquid culturing protocols to achieve higher starting cell densities, our overall success rate has climbed substantially. Second, although the protocols we provide can in principle be run with very little specific prior training or programming experience, some working knowledge of Python programming in general, and the Opentrons Python API in particular, is helpful. And third, the logistical challenges of moving from hundreds to potentially tens of thousands of samples—including storage, labeling, and in particular sample provenance validation and metadata tracking—are largely unaddressed here. We will be continuing to address each of these issues in future development of these protocols.

To ensure the greatest utility of our workflow for the research community, all protocols and hardware schematics are freely available for public use at https://github.com/tanaes/Moeller_Opentrons_protocol_library, https://github.com/tanaes/opentrons_functions, and https://github.com/CUMoellerLab/Labware. These repositories will be maintained and updated as we make further additions and improvements to the protocols in the future.

The isolates sequenced in this study represent, to our knowledge, the first large-scale compendium of cultured genomic resources from wild chimpanzee gut microbiomes. They were cultivated and sequenced by a single technician / postdoctoral research team over the course of just a few months, using samples that were not collected with bacterial isolation in mind, and were stored for over a decade at −80 °C in RNALater. The genome resources generated from this chimpanzee gut bacterial isolate collection complement and enable comparative analyses with existing gut bacterial genome databases derived from humans. Moreover, all isolates generated by this study have been preserved in glycerol stocks and are available upon request for research purposes.

The vast majority of global microbial genomic diversity remains unexplored. While centralized efforts to explore microbial diversity of particular significance to human health or economics are generating enormous amounts of new data, exploration of most other environments most often occurs in a more decentralized fashion, often by researchers with less access to the capital equipment and economies of scale enjoyed by their medically-oriented peers. Our intention in developing these protocols is to make high-throughput microbial genomics more accessible to more researchers, thereby increasing the diversity of environments from which microbial isolates and reference genomes can be obtained. Given the interconnectedness of microbial genomic diversity in nature, expanding the breadth of such data will be of substantial benefit to researchers studying microbes from all sorts of environments.

## Availability of data and materials

All raw sequence data from this publication are available in the Qiita data repository, study number 14410 (https://qiita.ucsd.edu/study/description/14410), as well as at the EBI ENA repository with accession number ERP136830. Opentrons protocols are available at https://github.com/CUMoellerLab/Moeller_Opentrons_protocol_library and custom function library at https://github.com/CUMoellerLab/opentrons_functions. Printable labware files and assembly instructions are available at https://github.com/CUMoellerLab/Labware. Isolates are available upon request.

## Funding

This work was supported by a laboratory start-up grant from Cornell University to AHM and grant R35 GM138284 from the National Institute of General Medical Sciences to AHM. JGS was partially supported by grant T32 AI145821 to the Cornell Institute for Host–Microbe Interaction and Disease by the National Institutes of Health.

## Authors’ Contributions

JGS and AHM conceived of the project and wrote the manuscript. WY and JGS performed laboratory work. JGS designed apparatus, wrote software and protocols, and performed analysis.

## Acknowledgements

We would like to thank Ilana Brito and all the members of the Moeller Lab for helpful discussions; as well as the Ithaca Generator makerspace, for contributing tools and expertise that aided in the design and production of the open-source labware in this study.

## Competing interests

JGS is the founder and sole proprietor of Lightweight Labware LLC. All other authors declare no competing interests.

## Supplementary Information

*Supplementary Figures*

**Figure S1. Alluvial plot of protocol efficiency.** Results from initial rounds of screening, showing samples passing certain QC thresholds at each stage (extraction: 0.1 ng/μL DNA concentration; Library prep: 0.5 ng/μL DNA concentration; Sequencing: 25 Mbp sequence yield; Taxonomy: taxonomy assigned by Sourmash; Assembly: High, Medium, and Low-quality assemblies), Contaminated assemblies, and unassembled samples. Colored lines connect the same sample through each stage of the chart.

**Figure S2. Relationship between assembly quality and library concentration**. Kernel density plots showing distribution of sequence library DNA concentrations for each level of assembly quality. As expected, assembly quality generally increases with library concentration, with libraries above 1 ng/μL typically yielding medium or high quality assemblies. Contaminated assemblies had library concentrations very similar to “high quality” uncontaminated libraries, suggesting contamination from multiple inocula rather than low-concentration reagent contamination.

**Figure S3. Intraspecific nucleotide diversity.** Pairwise Average Nucleotide Diversity among strains within each cluster show different patterns of within-‘species’ diversity revealed by whole-genome screening. Heatmap color values indicate log pairwise nucleotide diversity between each pair of isolates in a cluster. Color bars at left and annotations at right show host species identity of the sample from which the isolate was recovered. Color bars at top show the host individual. Different patterns highlight differences in the distribution and quantity of nucleotide variation in different isolate clusters, ranging from nearly clonal isolates recovered from within a single (clusters 30 and 34) or across multiple (cluster 41) individuals; to moderate amounts of variation largely partitioned across host individuals (cluster 16); to highly structured variation possibly indicative of multiple ‘species’ grouping within the same 95% ANI threshold (cluster 7 and 36).

**Figure S4. Genome-wide recombination rates.** 50kbp sliding window plots of recombination events inferred by ClonalFrameML for each 95% ANI strain cluster recovered in our analysis. All plots share the same Y axis scale, with a minimum of 0 and a maximum of 30 inferred recombination events per 50 kbp of genome length per genome. All plots have independent X axis scales, representing the total length of the assembly of the reference genome for that isolate cluster. Substantial variation in inferred recombination is apparent both among and within isolate genomes.

**Figure S5. Elevated copy number of putative plasmids.** Scatter plot shows the approximate coverage of putative plasmids in each assembly against the mean approximate coverage of contigs for that assembly. Orange dotted line denotes equal coverage. Coverage estimates were derived from SKESA.

*Supplementary Tables*

**Table S1: Sample information**

**Table S2: Culturing information**

**Table S3: Cost estimates**

**Additional File 1: Isolate taxonomic information, genome assembly statistics, and other metadata**

