## Supplemental Information for "Low-cost genomics enable high-throughput isolate screening and strain-level microbiome profiling"

### Supplementary Information

#### Supplementary Figures

##### Figure S1. Alluvial plot of protocol efficiency.

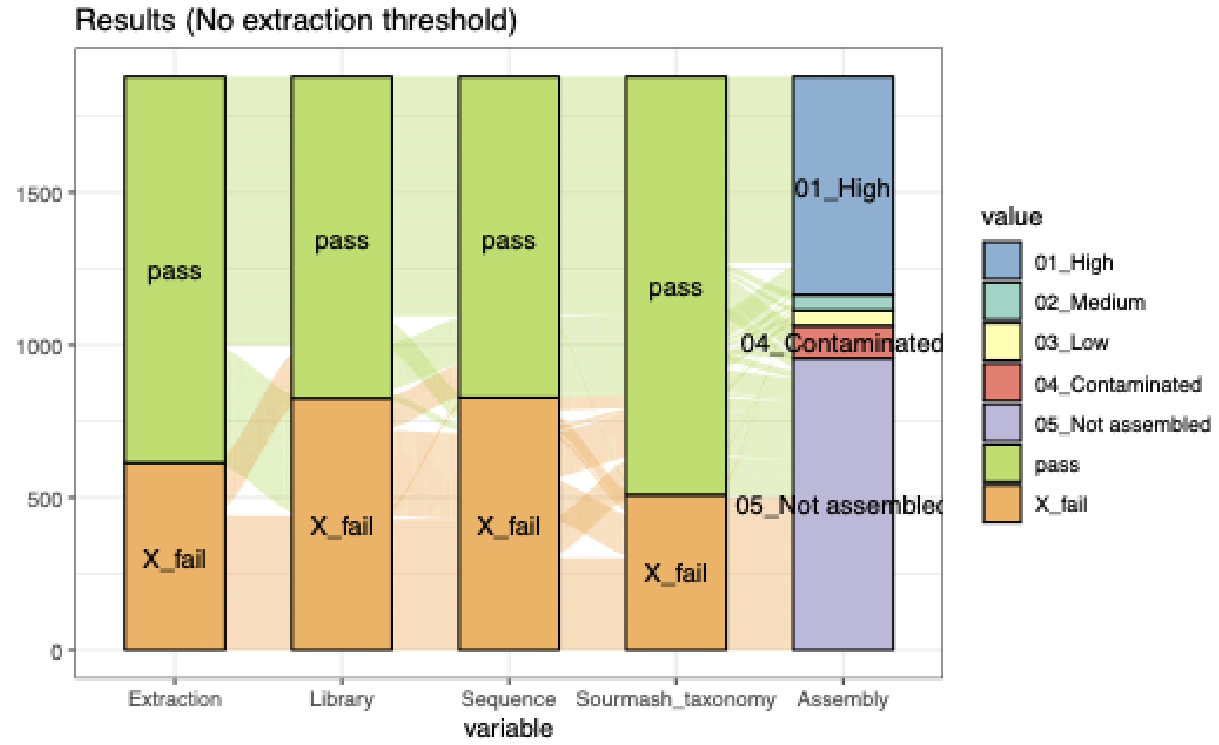

**Figure S1. Alluvial plot of protocol efficiency.** Results from initial rounds of screening, showing samples passing certain QC thresholds at each stage (extraction: 0.1 ng/µL DNA concentration; Library prep: 0.5 ng/µL DNA concentration; Sequencing: 25 Mbp sequence yield; Taxonomy: taxonomy assigned by Sourmash; Assembly: High, Medium, and Low-quality assemblies), Contaminated assemblies, and unassembled samples. Colored lines connect the same sample through each stage of the chart.

Figure S2. Relationship between assembly quality and library concentration.

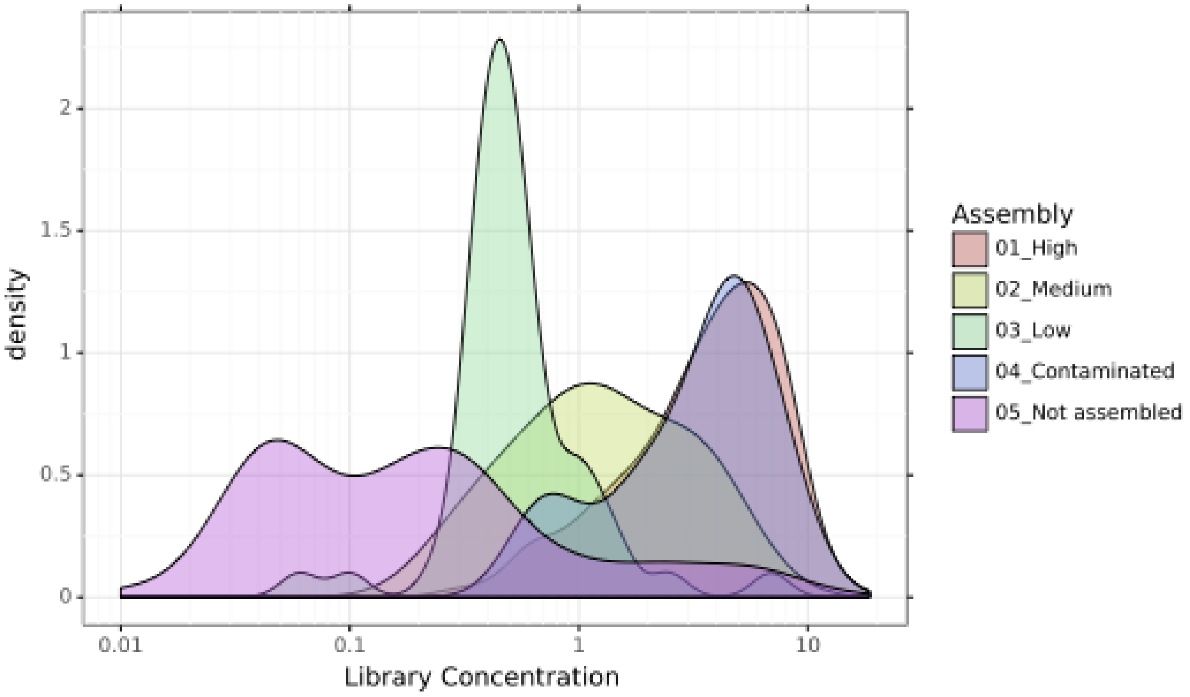

**Figure S2. Relationship between assembly quality and library concentration**. Kernel density plots showing distribution of sequence library DNA concentrations for each level of assembly quality. As expected, assembly quality generally increases with library concentration, with libraries above 1 ng/µL typically yielding medium or high quality assemblies. Contaminated assemblies had library concentrations very similar to "high quality" uncontaminated libraries, suggesting contamination from multiple inocula rather than low-concentration reagent contamination.

### Figure S3. Intraspecific nucleotide diversity.

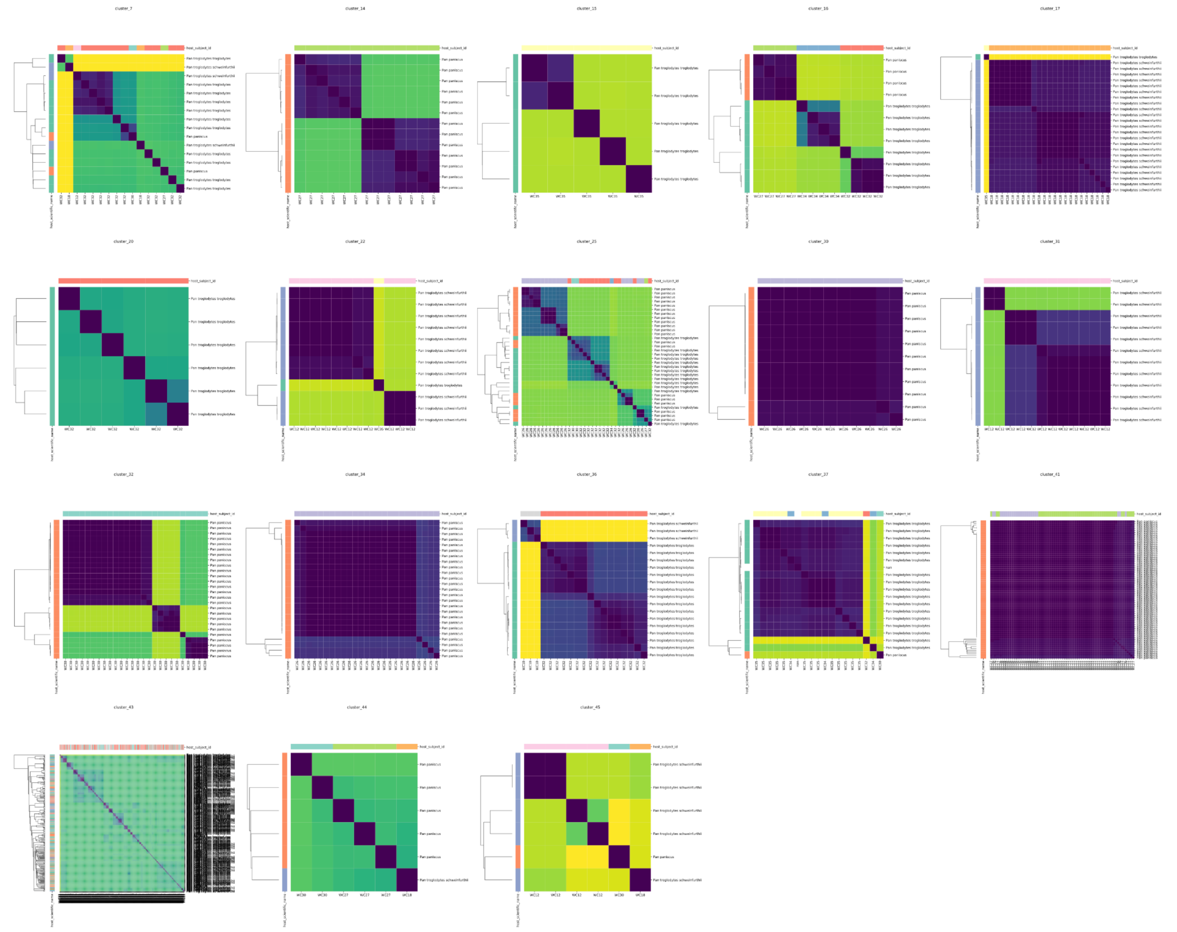

**Figure S3. Intraspecific nucleotide diversity.** Pairwise Average Nucleotide Diversity among strains within each cluster show different patterns of within-‘species’ diversity revealed by whole-genome screening. Heatmap color values indicate log pairwise nucleotide diversity between each pair of isolates in a cluster. Color bars at left and annotations at right show host species identity of the sample from which the isolate was recovered. Color bars at top show the host individual. Different patterns highlight differences in the distribution and quantity of nucleotide variation in different isolate clusters, ranging from nearly clonal isolates recovered from within a single (clusters 30 and 34) or across multiple (cluster 41) individuals; to moderate amounts of variation largely partitioned across host individuals (cluster 16); to highly structured variation possibly indicative of multiple ‘species’ grouping within the same 95% ANI threshold (cluster 7 and 36).

Figure S4. Genome-wide recombination rates.

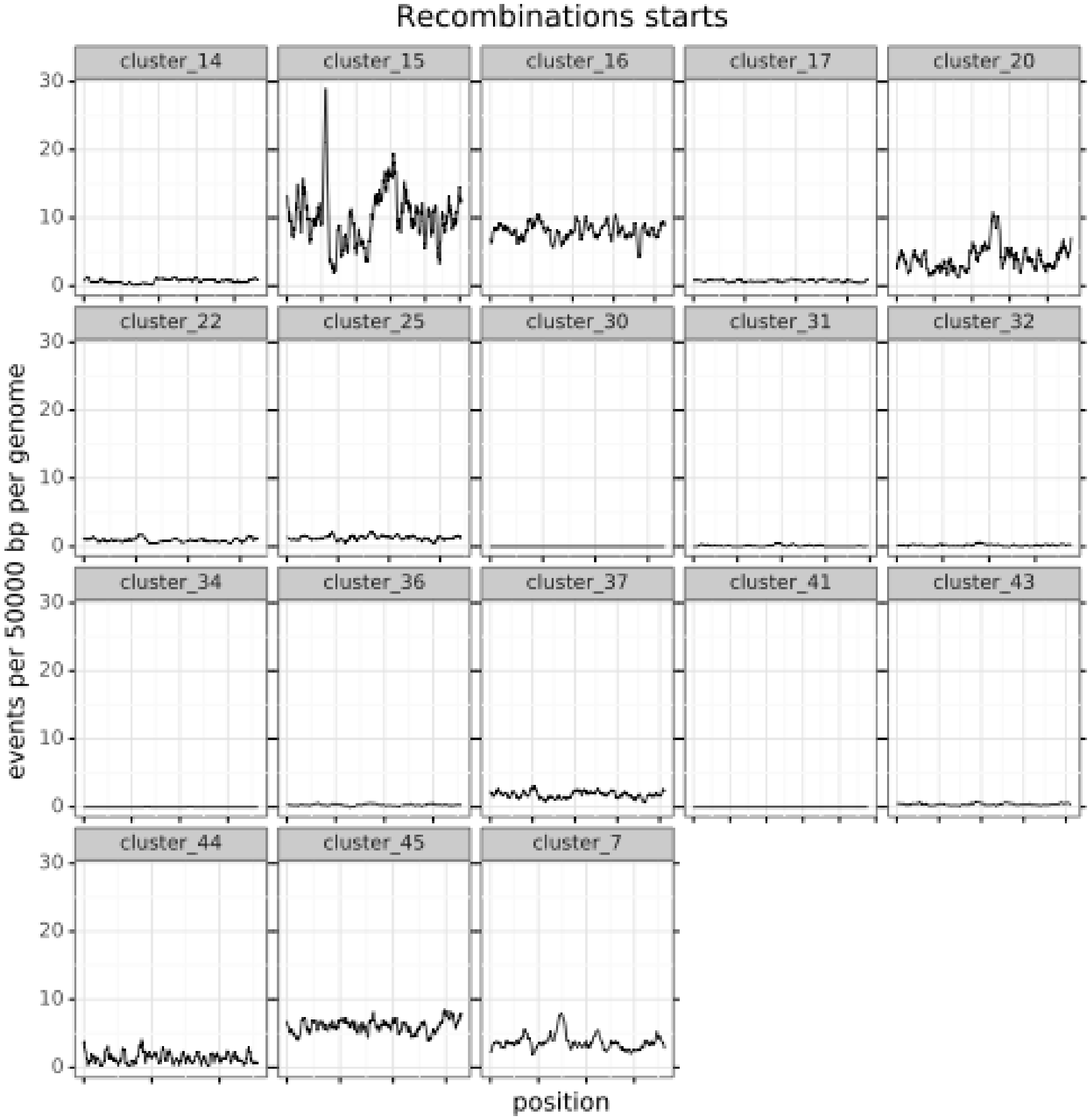

**Figure S4. Genome-wide recombination rates.** 50kbp sliding window plots of recombination events inferred by ClonalFrameML for each 95% ANI strain cluster recovered in our analysis. All plots share the same Y axis scale, with a minimum of 0 and a maximum of 30 inferred recombination events per 50 kbp of genome length per genome. All plots have independent X axis scales, representing the total length of the assembly of the reference genome for that isolate cluster. Substantial variation in inferred recombination is apparent both among and within isolate genomes.

Figure S5. Elevated copy number of putative plasmids.

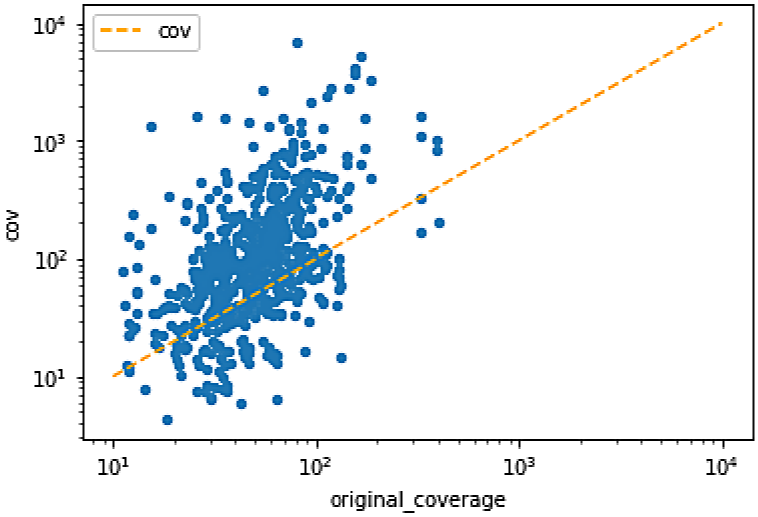

**Figure S5. Elevated copy number of putative plasmids.** Scatter plot shows the approximate coverage of putative plasmids in each assembly against the mean approximate coverage of contigs for that assembly. Orange dotted line denotes equal coverage. Coverage estimates were derived from SKESA.

### Supplementary Tables

#### Table S1: Sample information

| Individual | GM_Code | Collection_Date | Name | Sex | Subspecies | Site |
|---|---|---|---|---|---|---|
| WC02 | GM0566 | 2004-08-30 | Ch-095 | *F* | *Pan troglodytes schweinfurthii* | KL |
| WC10 | GM3878 | 2014-08-17 | Kati (Tita) | *F* | *Pan troglodytes schweinfurthii* | KL |
| WC12 | GM3909 | 2014-08-27 | Kazi | *M* | *Pan troglodytes schweinfurthii* | KL |
| WC18 | GM3902 | 2014-08-26 | Pairotti_Poa | *F* | *Pan troglodytes schweinfurthii* | KL |
| WC26 | KSG2915 | 2010-05-18 | IK2915 | | *Pan paniscus* | Ikela, Bafeke-Balanga |
| WC27 | KSG3821 | 2012-11-19 | TL3821 | | *Pan paniscus* | Tshuapa-Lomami-Lualaba |
| WC30 | KSG3845 | 2012-11-26 | TL3845 | | *Pan paniscus* | Tshuapa-Lomami-Lualaba |
| WC32 | DP130 | 2003-11-07 | MP130 | | *Pan troglodytes troglodytes* | Cameroon, Doumo Pierre |
| WC34 | DP095 | 2003-07-29 | MP95 | | *Pan troglodytes troglodytes* | Cameroon, Doumo Pierre |
| WC35 | GT504 | 2005-02-22 | | | *Pan troglodytes troglodytes* | RC, Goualougo Triangle |

### Table S2: Culturing information

| Plate | Host | Culture date | Culture medium |
|---|---|---|---|
| WC10-BSM-1 | WC10 | 2020-2-17 | BSM |
| YS12 | WC12 | 2020-2-27 | YCFA+Starch |
| BBE18 | WC18 | 2020-2-27 | BBE |
| BSM18-1 | WC18 | 2020-2-27 | BSM |
| BSM18-2 | WC18 | 2020-2-27 | BSM |
| YS18 | WC18 | 2020-2-27 | YCFA+Starch |
| YS-26-1 | WC26 | 2020-2-20 | YCFA+Starch |
| YS-26-2 | WC26 | 2020-2-20 | YCFA+Starch |
| 27-YS-1 | WC27 | 2020-2-20 | YCFA+Starch |
| 27-YS-2 | WC27 | 2020-2-20 | YCFA+Starch |
| BHIS-30 | WC30 | 2020-2-17 | BHIS |
| WC-30 | WC30 | 2020-2-17 | YCFA |
| YS-30-3 | WC30 | 2020-2-17 | YCFA+Starch |
| YS-30 | WC30 | 2020-2-17 | YCFA+Starch |
| 32-YCFA-2 | WC32 | 2020-2-17 | YCFA |
| BHIS-32 | WC32 | 2020-2-17 | BHIS |
| BSM-32 | WC32 | 2020-2-17 | BSM |
| WC-32 | WC32 | 2020-2-17 | BHIS |
| WC32-BSM-2 | WC32 | 2020-2-17 | BSM |
| BBE-34-35 | WC34 | 2020-2-24 | BBE |
| YS-35-1 | WC35 | 2020-2-24 | YCFA+Starch |
| YS-35-2 | WC35 | not recorded | YCFA+Starch |

### Table S3: Cost estimates

| Item | Step | Qty used[1] | Stock price | Stock qty | Stock unit | Net cost |
|---|---|---|---|---|---|---|
| 96 well strip plate | Extraction | 1 | $269.90 | 50 | ea | $5.40 |
| GITC buffer | Extraction | 40 | $28.30[2] | 1000 | mL | $1.13 |
| Magnetic beads | Extraction | 2.5 | $223.00 | 24 | mL | $23.23 |
| Isopropanol | Extraction | 30 | $19.05 | 4000 | mL | $0.14 |
| Ethanol | Extraction | 60 | $25.00 | 4000 | mL | $0.38 |
| VWR deep well plate | Extraction | 1 | $187.85 | 50 | ea | $3.76 |
| Bio-Rad PCR plate | Extraction | 1 | $1,366.40 | 400 | ea | $3.42 |
| OpenTrons 300 µL tips | Extraction | 2 | $3,025.00 | 1000 | ea | $6.05 |
| OpenTrons 200µL filter tips | Extraction | 2 | $687.50 | 100 | ea | $13.75 |
| glass lysis beads | Extraction | 48 | $40.00 | 453 | g | $4.24 |
| | | | | | **Subtotal:** | **$61.49** |
| Illumina Library Prep Kit | Library Prep | 96 | $1038.00 | 960 | rxn | $103.80 |
| Primestar GXL Polymerase | Library Prep | 225 | $671.00 | 1000 | units | $150.98 |
| Index primers | Library Prep | 1 | $500.00 | 16 | plates | $31.25 |
| SPRI beads | Library Prep | 6 | $3,835.80 | 500 | mL | $46.03 |
| Tagmentation buffer | Library Prep | 3.2 | $40.34[2] | 1000 | mL | $0.13 |
| Stop buffer | Library Prep | 1.6 | $0.86[2] | 1000 | mL | $0.00 |
| Wash buffer | Library Prep | 24 | $8.14[2] | 1000 | mL | $0.20 |
| Nuclease free H2O | Library Prep | 8 | $30.22 | 1000 | mL | $0.24 |
| Ethanol | Library Prep | 36 | $25.00 | 4000 | mL | $0.23 |
| Bio-Rad PCR plate | Library Prep | 3 | $1,366.40 | 400 | ea | $10.25 |
| OpenTrons 300 µL tips | Library Prep | 1 | $3,025.00 | 1000 | ea | $3.03 |
| OpenTrons 200µL filter tips | Library Prep | 1 | $687.50 | 100 | ea | $6.88 |
| OpenTrons 20 µL tips | Library Prep | 1 | $3,025.00 | 1000 | ea | $3.03 |
| OpenTrons 20 µL filter tips | Library Prep | 2 | $687.50 | 100 | ea | $13.75 |
| | | | | | **Subtotal:** | **$369.77** |
| NextSeq 550 | Sequencing | 11 | $5270.00 | 110 | Gbp | $527.00 |
| | | | | | **Subtotal:** | **$527.00** |
| **Per-sample costs** | | | | | | |
| Extraction | $0.64 | | | | | |
| Library Prep | $3.85 | | | | | |
| Sequencing | $5.49 | | | | | |
| **Total** | **$9.98** | | | | | |

[1] Per 96-well plate
[2] Home-made buffer, ‘stock price’ indicates cost of constituent materials

Additional File 1: Isolate taxonomic information, genome assembly statistics, and other metadata
